# Can tropical agamid lizards physiologically tolerate the altered thermal mosaic of urban microhabitats?

**DOI:** 10.64898/2026.04.06.716463

**Authors:** Marwa Razak, Amanda Ben, Saumitra Dhere, Maria Thaker

## Abstract

Urbanization and human-induced environmental changes create unique and unprecedented thermal landscapes, yet the extent to which species respond to these changes remains poorly understood. One major challenge in studying these responses is the spatial mismatch between the small scale at which organisms experience their environment and the broader scale at which climate data are typically collected. We use Infrared Thermography (IRT) to quantify the fine scale microclimate in urban and rural habitats used by two tropical agamid lizards, *Calotes versicolor* and *Psammophilus dorsalis*. By combining field-based body temperatures and lab-based measures of thermal limits (CT_max_, CT_min_)and preferences (T_pref_), we assess how the thermal heterogeneity of these fine mosaics of microhabitats influence the degree of thermoregulation (k) of these species. We find that thermal responses to urbanization are shaped by species-specific thermal traits and patterns of microhabitat use. Between the species, urban individuals did not differ markedly in habitat thermal heterogeneity, substrate temperature used or degree of thermoconformity. However, within species, *P. dorsalis* experiences warmer and more heterogeneous conditions in rural habitats, whereas *C. versicolor* experiences similar thermal conditions across habitats. *Calotes versicolor* also exhibits broader thermal tolerance and preferred temperature ranges than *P. dorsalis*. Collectively, our results suggest that *P. dorsalis* may be more susceptible to the thermal constraints imposed by human-modified landscapes. Overall, we demonstrate the critical need to account for microclimatic conditions and species-specific thermal traits when determining how animals respond to changes in the thermal environment expected from climate change.

## 1. INTRODUCTION

Urbanisation and associated climate warming are accelerating at an alarming rate and is undeniably reshaping environments. There is now no doubt that many animals are already affected by and are responding to the present anthropogenically-driven changes to environments (Parmesan & Yohe, 2003; Walther et al., 2002). Ectotherms may be disproportionately susceptible to these effects as they rely on external heat sources to regulate their body temperature (Aragón et al., 2010). For ectotherms, environmental temperature is one of the most important factors that governs basic physiological functions such as growth, reproduction, and performance (Deutsch et al., 2008). Owing to their high thermal sensitivity (Huey & Kingsolver, 1989) and rapid response to changes in their environment such as shifts in distribution, abundance, and diversity, several ectotherms have been found to be valuable bioindicators of climate change (Chowdhury et al., 2023). Given that ectotherms make up majority of terrestrial biodiversity (Wilson, 1999), it is essential to examine how temperature affects these organisms. Determining these effects helps in making informed predictions about the future of biodiversity under the influence of increased urbanization and climate change (Amarasekare & Savage, 2012).

Predictive models to understand how species will respond to climate change are crucial and have gained significant attention. Most of these models, however, are at the macroclimatic scale and typically assess the effects of changes in monthly, seasonal, or annual environmental conditions (Pincebourde et al., 2016). Organisms, on the other hand, do not experience climate on such broad spatial scales. Instead, they live in thermally heterogeneous microhabitats that may have diverse microclimates that are not reflected in measures of regional temperature (Kearney et al., 2009; Potter et al., 2013). For example, under same ambient temperatures, leaf surfaces of canopies exposed to the sun were found to be about 8 degrees warmer than that of canopies in the shade (Stoutjesdijk, 1977). The topography of the microhabitat can also create pronounced temperature differences at small spatial scales (Sears et al., 2011). For instance, a single rock can display a temperature difference of at least 9 degrees over a few centimeters (Pike et al., 2012). Ultimately, the microclimate experienced by an organism is determined by spatial and temporal variations in temperature within its habitat and the extent to which the organism can effectively utilize these (Pincebourde & Suppo, 2016). Consequently, increased variability in habitat provides a greater potential for more precise regulation of temperature, which in turn corresponds to better fitness of the organism (Gaudenti et al., 2021).

The significance of microclimate variation in shaping the thermal physiology of organisms has emerged as an important area of research (Pincebourde et al., 2016). Microclimatic variation within habitats can drive changes in thermal traits comparable to those observed across broad geographic temperature gradients (Pincebourde & Woods, 2020). Key physiological traits such as thermal tolerance (CT_max_, CT_min_) and preferred body temperature (T_pref_) are important measures of thermal responses (Caldwell et al., 2015; Duarte et al., 2012). Variation in microhabitat use has been shown to significantly influence these traits. For instance, in sympatric species of army ants, above-ground species exposed to higher surface temperatures exhibit significantly higher critical thermal maxima (CT_max_) than below-ground species (Baudier et al., 2015). Similarly, in Aegean wall lizards, color morphs that occupy different microhabitats show differences in preferred body temperatures (T_pref_), with individuals from cooler microhabitats showing lower T_pref_ values. Thus, the availability of fine-scale thermal heterogeneity within microhabitats can be important for behavioral thermoregulation and can influence thermal traits (Kearney et al., 2009).

While thermal traits have been widely used to assess the responses of species to global climate change, this approach is also relevant in the context of urban warming (Diamond et al., 2017). Urban environments typically exhibit higher average temperatures than rural areas, creating localized warming known as urban heat islands (Battles & Kolbe, 2019; Vardi et al., 2023). Urban microclimates therefore offer analogs for future climatic conditions and serve as natural laboratories for forecasting ecological responses to warming (Youngsteadt et al., 2015). Examining the climate exposure within these microhabitats relative to the thermal limits and preferences of species is crucial for identifying their vulnerability to changing temperatures (Scheffers et al., 2013). Along urban-rural gradients, significant intraspecific variation in thermal traits has been observed in some taxa. For instance, acorn ants from urban areas exhibit both higher heat tolerance and lower cold tolerance than those from rural populations (Diamond et al., 2018). Similar trends were observed in leaf-cutter ants (*Atta sexdens*), where urban ants show higher upper thermal tolerance (CT_max_) compared to their rural counterparts (Angilletta et al., 2007). Such differences in thermal traits between urban and rural populations, however, are not obligate. For example, urbanisation seems to have no effect on the preferred temperature (T_pref_) of the ground-dwelling spider species, *Pardosa saltans* (Cabon et al., 2023). A lack of association may be because spiders can exploit the thermal heterogeneity within their microhabitats to maintain optimal body temperatures, thereby weakening the selection pressure on thermal physiology (Bogert effect; Huey et al., 2003). It is also possible for studies like these that measurements of ambient habitat temperatures are not, in fact, reflective of the actual microclimatic conditions experienced by the animals.

Here we use Infrared Thermography (IRT) to quantify fine scale microclimatic differences of urban and rural microhabitats of two co-occurring agamid lizards with differing microhabitat preferences. The common garden lizard, *Calotes versicolor*, is a geographically widespread semi-arboreal species that is commonly found in open scrubland with low vegetation and deciduous forests (Wang et al., 2023). In urban habitats, they are often seen in open areas with low vegetation, perched on rocky pole fences. The peninsula rock agama, *Psammophilus dorsalis*, is endemic to India, with a broad distribution across open rocky terrain interspersed with boulders and scrubs (Chandra & Gajbe, 2005; Daniel, 2002). In urban environments, *P. dorsalis* is frequently observed on wall ledges, rocks, and bricks (Balakrishna et al., 2021). We sampled lizards from both urban and rural populations to investigate how thermal traits, specifically preferred body temperature (T_pref_) and critical thermal limits (CT_max_, CT_min_), vary between the species and across different habitats. We coupled this information with field measures of body temperature and microhabitat temperature, using Infrared Thermography (IRT). We predict that urbanisation would lead to thermal homogenisation of microhabitats, reducing the range of temperatures available for effective thermoregulation. Further, we expect that the thermal physiology of these species would differ between the urban and rural populations, in ways that reflect the differences of their thermal environments. By accounting for both realised microhabitat temperatures and organismal thermal capacities, our approach allows us to better understand how urbanisation influences the responses of vertebrate ectotherms in the tropics. We integrated detailed measurements of local thermal environments, behavior, and physiology to test whether two closely related species of tropical ectotherm are likely to respond to climate change in the same way.

## 2. METHODS

### 2.1 Study sites

We sampled several sites across three districts of Karnataka, India - Bengaluru, Kolar, and Ramanagara, where both species occur. Bengaluru, located in southern peninsular India, is one of the country’s five fastest-growing cities, shaped by rapid urban development and economic growth. Elevationally, Bengaluru (12°59′ N, 77°57′ E) lies about 920 m a.s.l, and receives a mean annual rainfall of 880 mm (Ramachandra & Kumar, 2010). On the east, Kolar (13°08′N 78°08′E) lies at 822 m a.s.l with characteristic semi-arid land and hard rocky terrain and receives a mean annual rainfall of 744 mm (Kumara, 2018). Southwest of Bengaluru, Ramanagara (12.72° N, 77.27° E) lies at 742.5 m a.s.l with a forest cover of 17.21% of its total area and receives an average rainfall of 931.58 mm annually (Nagaraja et al., 2025). Lizards were collected from urban (n = 5 sites) and rural (n = 5 sites) areas, which were at least ca. 10km from each other. Urban sites were within residential areas with undeveloped land, cement block walls, scrub vegetation, and frequent human disturbance. Rural sites were relatively undisturbed and comprised of rocky outcrops and boulders interspersed with scrub vegetation for *P. dorsalis*; and open scrubland and low vegetation open forests (including invasive *Lantana camara*) for *C. versicolor*.

### 2.2 Animal housing

All data were collected between May 2023 to July 2024, during the active season for these species. Adult lizards were caught using a telescopic fishing pole with a lasso attached to the end. Following capture, lizards were transferred to individual cloth bags and transported to the laboratory in cooler boxes to reduce thermal stress during transport. At the laboratory, lizards were housed individually in glass terraria (60 × 30 × 25 cm) in a dedicated animal housing room maintained at 25-27°C with a 12-hour photoperiod (0700-1900 h). Terraria were lined with disposable paper towels as substratum and provided with rocks for basking, and a plastic pipe for refuge. Incandescent lights (60 W) above each tank were turned three times daily for 30 min to provide basking opportunities. Lizards were fed with mealworms daily and provided with water ad libitum in petri dishes. Before experiments, mass (g) and snout-to-vent length (SVL, mm) of all individuals were measured. Lizards were given 1 day to habituate to the laboratory conditions, before T_pref_ was measured. After a 24-hour recovery period, lizards were either used for CT_max_ or CT_min_ measurements. To prevent physiological adaptation to laboratory settings, all thermal data were collected within two weeks of capture, after which lizards were released at their capture sites.

### 2.3 Preferred Temperature (T_pref_)

Preferred temperature (T_pref_) data were obtained from 49 adults of *C. versicolor (*n = 26 rural, 23 urban) and 119 adults of *P. dorsalis* (n = 68 rural, 54 urban). A thermal gradient (200 cm long x 12.5 cm high x 40 cm wide) was constructed with a wooden frame separated into four lanes by opaque plastic separations, with the base lined with sandpaper to provide traction for lizards. Four infrared heating lamps were positioned above on one end of each lane and cool ice packs placed on the opposite ends, resulting in a thermal gradient ranging from 8 - 52 °C (Figure S1). Lizard body temperature was measured using thermocouple probes (Amprobe TMD-50 Thermocouple) inserted into the cloaca and fastened by adhesive medical tape. Lizards were introduced to the middle of each lane of the thermal gradient and were allowed to habituate for an hour. Body temperature was recorded every 10-minutes over the following 2-hour period, and we calculated T_pref_ as the median value of the 14 cloacal readings.

### 2.4 Critical thermal limits (CT_max_, CT_min_)

Critical thermal limit data were collected for both *P. dorsalis* (n = 67 rural, 53 urban) and *C. versicolor (*n = 26 rural, 22 urban). We measured CT_min_ and CT_max_ as the temperature at which the lizard loses locomotor functions following (Clifton et al., 2021). To monitor body temperature during trials, a thermocouple probe (Amprobe TMD-50 Thermocouple) was inserted into the cloaca and fastened with medical tapes. For CT_max_, each lizard was placed in a steel container submerged in an electric water bath in which the temperature was gradually increased at a rate of 1°C/min. Once the lizard looked visibly sluggish, it was flipped onto its back and tested for the ability to right itself at every 1 minute. The method used to determine CT_min_ was similar by decreasing the temperature at 1 °C/min by adjusting the position of ice around the container in an ice-bath. These values were also used to determine the thermal tolerance range, calculated as the mean difference between CT_max_ and CT_min_ following Calosi et al., (2008).

### 2.5 Thermal imaging of microhabitats in the wild

Thermal imaging of microhabitats involved capturing and marking lizards, followed by thermal imaging over subsequent days. We sampled only adult lizards with snout-to-vent length (SVL) > 70 mm to reduce variation due to body size. Following capture, we measured SVL using a ruler and body mass using a portable weighing scale. Lizards were marked with a nontoxic permanent marker (POSCA) and released at their site of capture. The same site was revisited on subsequent days and visually scanned to locate marked individuals. We took thermal images of the individuals and the substrates they were perched on from about 4 - 9 m distance away using a FLIR T640 thermal camera (Goller et al., 2014). Thermal images were taken every 2 hours during the entire period of activity ranging from 0800-1800 hours (see Figure S2).

Immediately after imaging lizards, we imaged four unused substrates surrounding the lizard at angles of 90°, 180°, and 270° to assess available thermal niches similar to Mohanty et al., (2021).

Thermal images were processed using FLIR Thermal Studio Pro software. From these images, which comprise of 307,200 (640 x 480) pixels each, we excluded pixels with extreme temperature values, such as the sky. We used the draw tool function to measure (see Figure 1) - (a) mean body temperature along the dorsal line from head to tail base (Goller et al., 2014); (b) substrate temperature (area adjacent to the lizard, measured as a rectangle proportional to the line drawn), and (c) mean environment temperature from substrates around the sampled lizard (Mohanty et al., 2021). We used an R package, “ThermStats”(Senior et al., 2019) to quantify thermal heterogeneity by calculating Shannon Diversity Index (SHDI) and Simpson Diversity Index (SIDI) from each image.

**Figure 1.**
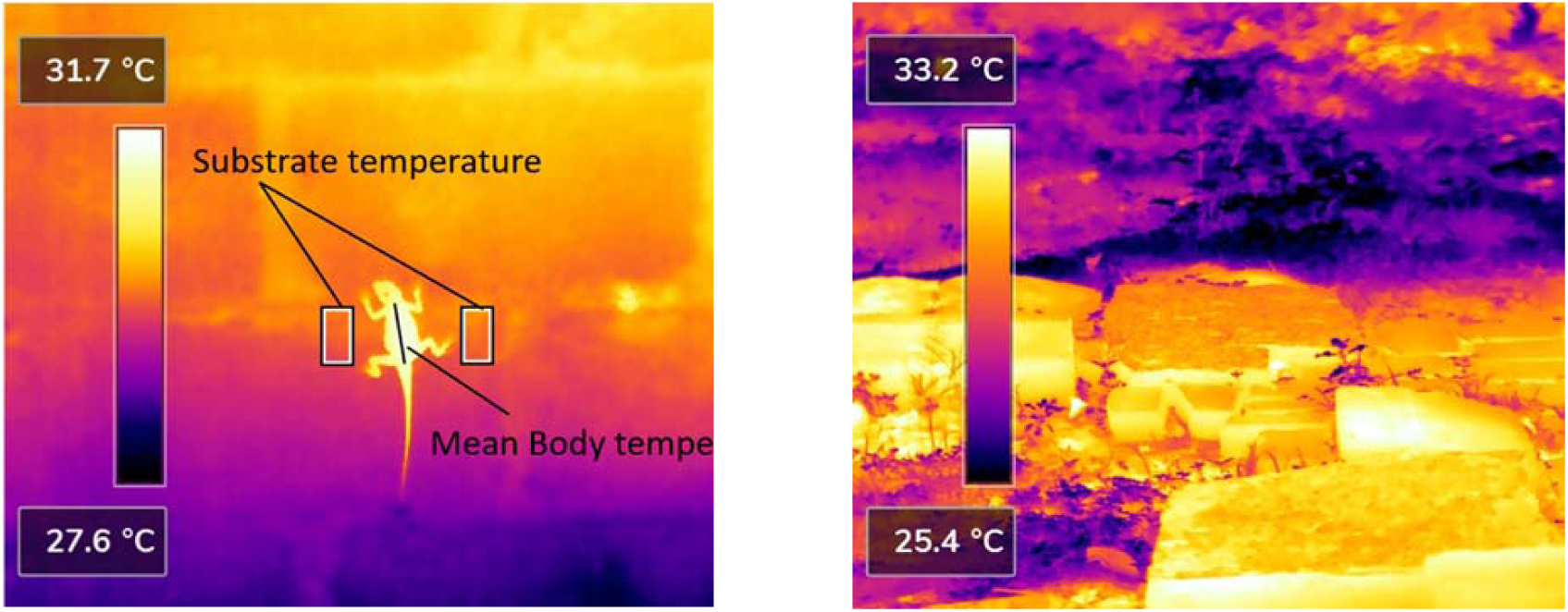
Thermal image from an urban site of *P. dorsalis*. (a) Mean body temperature of the lizard was calculated from a single line drawn on the dorsal side. Substrate temperature was measured from two regions next to the lizard (size of each region was proportional to the size of the lizard, excluding limbs and the tail). (b) Example of microhabitats around a lizard from which thermal heterogeneity (SHDI, SIDI) was calculated.

### 2.6 Statistical analysis

All statistical analyses were done using R studio. Normality of the data was assessed through Shapiro-Wilke’s tests, Q-Q plots, and histograms, and accordingly, parametric or nonparametric statistical tests were used.

To test differences in thermal profile measures (T_pref_, CT_max_ and CT_min_), we ran separate Generalised Linear Models (GLM) with SVL, habitat type (rural, urban), species (*C. versicolor*, *P. dorsalis*), and a habitat type by species interaction as predictors. Data for both sexes were pooled together as SVL is a good proxy for sex differences (males being larger in both species, see Table S1). GLMs were followed by Tukey’s post hoc tests, whenever relevant. For thermal tolerance range (CT_max_ - CT_min_), a t-test was used to assess differences between the species.

To test differences in microhabitat thermal heterogeneity, we ran GLMs with habitat type (rural, urban) and species (*C. versicolor*, *P. dorsalis*) as fixed effects and the diversity indices (SHDI, SIDI) as the response variable. Similar GLM models were used to assess the differences in mean substrate and body temperature from thermograms. Finally, to measure degree of thermoregulation (k), we ran a linear regression of body temperature and substrate temperature for urban and rural populations of *P. dorsalis* and *C. versicolor*. Based on the slope of this regression, k = 0 is considered a perfect thermoregulator and k = 1 a perfect thermoconformer as per (Huey & Slatkin, 1976).

## 3. RESULTS

### 3.1 Thermal traits

T_pref_ was significantly different between urban and rural lizards (Figure 2a, Table 1). Both *C. versicolor* and *P. dorsalis* from urban habitats had significantly higher T_pref_ values than rural ones (t = 3.14, p < 0.01). We found a significant difference between species (t = -4.10, p < 0.001) with *P. dorsalis* preferring relatively lower temperatures, but no significant interaction effect between habitat and species was observed (t = -0.01, p = 0.99). Furthermore, SVL did not significantly affect T_pref_ (t = -0.69, p = 0.49).

**Figure 2.**
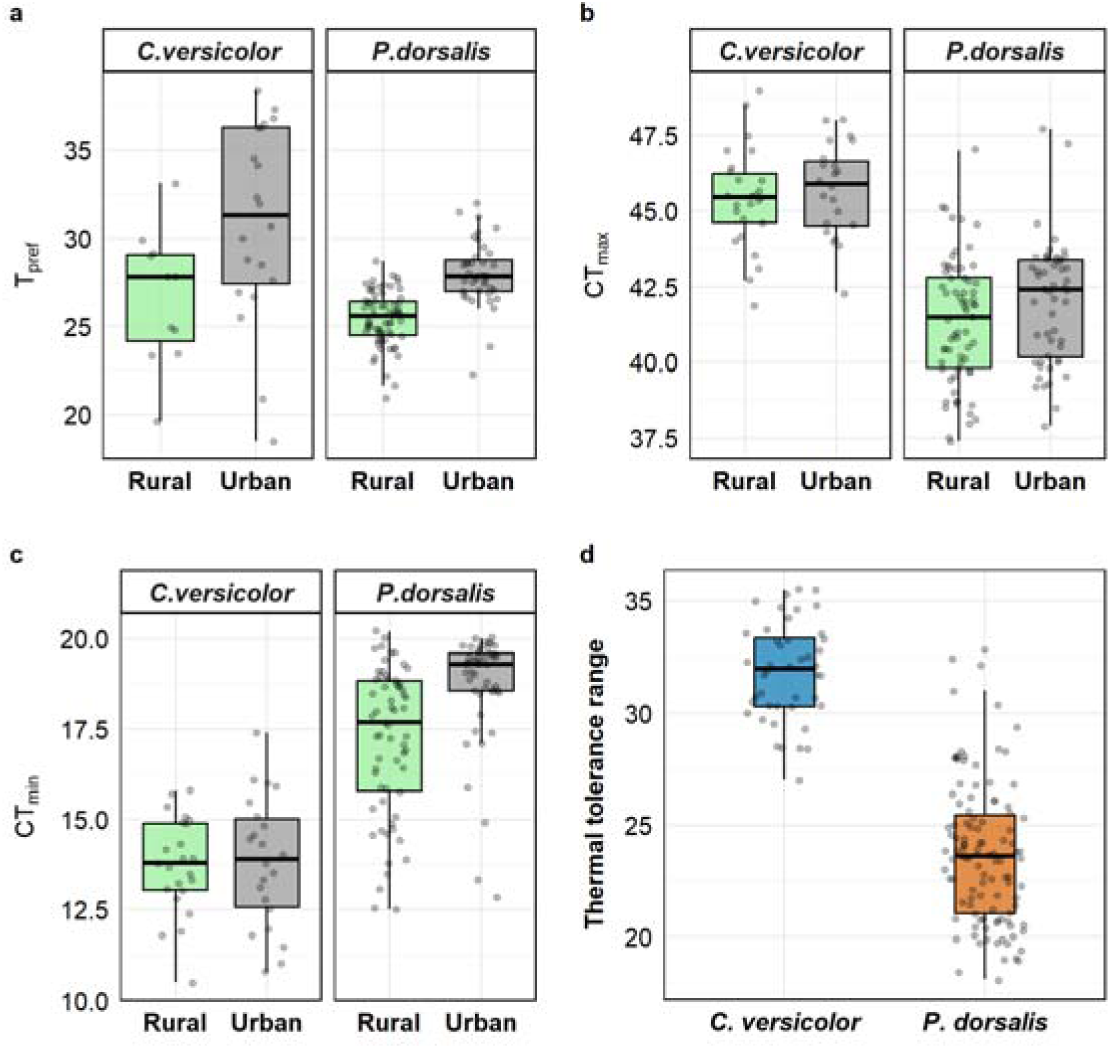
Thermal Profile of *C. versicolor* and *P. dorsalis* from rural (green) and urban (grey) habitats. a) Preferred Temperature (T_pref_), b) Critical Thermal Maxima (CT_max_), c) Critical Thermal Minima (CT_min_) and, d) Thermal tolerance range calculated as the mean difference between CT_max_ and CT_min_ for *C. versicolor (*yellow) and *P. dorsalis*(orange).

**Table 1.** Results of a Generalized Linear Model (GLM) assessing the effect of species, body size (SVL), and habitat on Tpref, CTmax, and CTmin. Values are presented as median ± SD and interquartile range (25%–75%) for each population. Values in bold denote statistical significance at p <0. 05.

|  | *C. versicolor* |  | *P. dorsalis* |  | Statistical Results |  |  |  |
| --- | --- | --- | --- | --- | --- | --- | --- | --- |
|  | Rural | Urban | Rural | Urban | Species | SVL | Habitat | Interaction |
| N | 26 | 21 | 67 | 39 |  |  |  | Species*Habitat |
| Tpref | 28.23 ± 3.77 | 30.70 ± 5.38 | 25.60 ± 1.54 | 27.85 ± 1.89 | **t = -4.1** | t = -0.69 | **t = 3.14** | t = -0.01 |
|  | 25.34–30.19 | 27.60–36.35 | 24.50–26.43 | 27.00–28.78 | **p < 0.001** | p = 0.49 | **p < 0.01** | p = 0.99 |
| N | 26 | 22 | 67 | 52 |  |  |  |  |
| CTmax | 45.45 ± 1.65 | 45.9 ± 1.5 | 41.5 ± 2.09 | 42.40 ± 2.01 | **t = -9.0** | **t = 2.07** | t = 0.63 | t = 0.24 |
|  | 44.63–46.23 | 44.53–46.65 | 39.8–42.8 | 40.18–43.4 | **p < 0.001** | **p = 0.04** | p = 0.53 | p = 0.81 |
| N | 23 | 22 | 63 | 48 |  |  |  |  |
| CTmin | 13.8 ± 1.34 | 13.9 ± 1.79) | 17.7 ± 2.03 | 19.3 ± 1.59 | **t = 7.96** | t = 0.3 | t = 0.17 | **t = 2.16** |
|  | 13.05–14.9 | 12.58–15.03 | 15.8–18.85 | 18.57–19.6 | **p < 0.001** | p = 0.76 | p = 0.87 | **p = 0.03** |

CT_max_ values differed significantly between the species, such that *C. versicolor* had higher CT_max_ than *P. dorsalis* (t = -9.0, p < 0.001; Figure 2b; Table 1). CT_max_ was not significantly different for lizards between urban and rural habitats (t = 0.63, p = 0.53), and there was no significant interaction between habitat type and species in predicting CT_max_ (t = 0.24, p = 0.81). SVL had a significant positive effect on CT_max_ overall (t = 2.07, p = 0.04).

For CT_min_, we found a significant interaction between habitat type and species (t = 2.16, p = 0.03; Figure 2c; Table 1). SVL did not significantly affect CT_min_ (t = 0.3, p = 0.76). Post hoc analysis showed that lizards from both habitats of *C. versicolor* had significantly lower CT_min_ relative to the corresponding populations of *P. dorsalis*. Within species, *P. dorsalis* from rural habitats had lower CT_min_ values than urban ones (t = -4.18, p < 0.0001). There was no significant difference in CT_min_ values across habitats for *C. versicolor (*t = -0.17, p = 0.87).

Overall, *C. versicolor* has a significantly broader thermal tolerance range (CT_max_ - CT_min_) than *P. dorsalis* (t = 18.14, p < 0.001, Figure 2d).

### 3.2 Thermal heterogeneity within microhabitats

We found a significant interaction between habitat and species for SHDI (t = −8.45, p < 0.001; Figure 3, Table S2). Thermal environment in rural habitats of *C. versicolor* had significantly lower SHDI values compared to those of *P. dorsalis* (t = −5.82, p < 0.001). In the urban habitats, SHDI values were higher for sites used by *C. versicolor* compared to those used by *P. dorsalis* (t = 6.51, p < 0.001). For *P. dorsalis*, SHDI values of rural sites were significantly higher than those of urban sites (t = 12.85, p < 0.001), while for *C. versicolor*, SHDI values was lower in rural habitats compared to urban ones (t = −3.66, p < 0.001).

**Figure 3.**
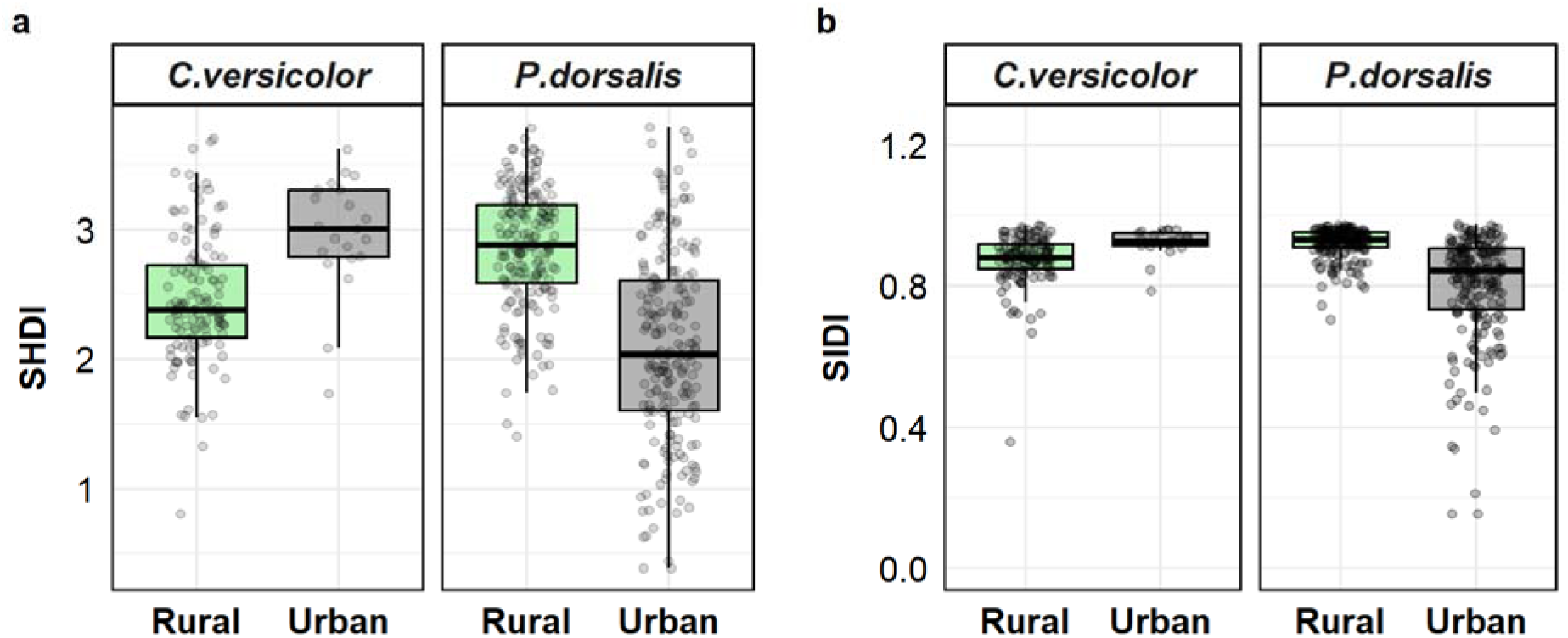
Thermal heterogeneity within the urban and rural microhabitats of *P. dorsalis* and *C. versicolor*, measured as (a) SHDI (Shannon Diversity Index) and (b) SIDI (Simpson Diversity Index) from thermal images.

For SIDI, we found a significant interaction effect between habitat and species (t =−6.383, p < 0.001; Figure 3; Table S2). Thermal microhabitats used by *C. versicolor* had a significantly lower SIDI compared to *P. dorsalis* in rural habitats (t = −3.8, p < 0.001). Urban environments had significantly higher SIDI for *C. versicolor* than *P. dorsalis* (t = 5.23, p < 0.001). Within species, rural habitats of *P. dorsalis* had significantly higher SIDI than urban ones (t = 11.58, p < 0.001), while there was no significant difference in SIDI between urban and rural habitats of *C. versicolor (*t = −1.96, p = 0.0503).

### 3.3 Body temperature and substrate temperature across habitats

Mean body temperature obtained from thermal images showed significant interaction between habitat and species (t = 4.98, p<0.001; Figure 4; Table S3). Rural lizards of *C. versicolor* had significantly lower body temperatures compared to rural *P. dorsalis* (t = -4.98, p < 0.001). However, species did not differ significantly in body temperature in the urban habitats (t = 1.32, p = 0.19). Within species, *P. dorsalis* had significantly higher body temperatures in rural than urban habitats (t = 6.44, p < 0.0001), while *C. versicolor* showed no significant difference across habitats (t = -1.39, p = 0.1649).

**Figure 4.**
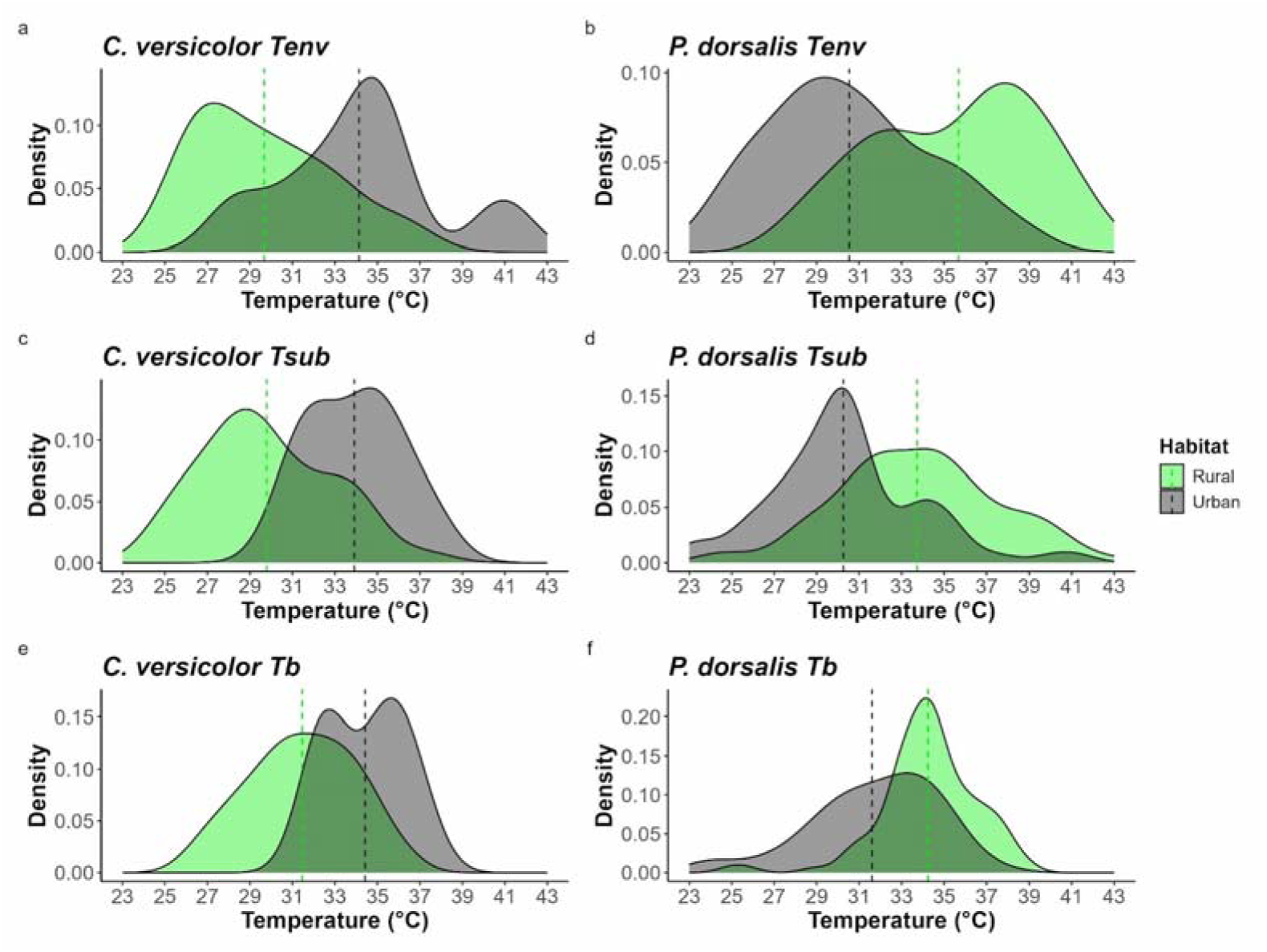
Density plots illustrating the frequency distribution of environmental temperature (Tenv), substrate temperature (T_sub_) and of body temperature (T_b_) for *C. versicolor (*left column: panels a, c, e) and *P. dorsalis* (right column: panels b, d, f) across rural (green) and urban (grey) habitats, measured from thermal images. The vertical dashed line represents mean temperature for lizards from each habitat type.

Similarly, mean substrate temperature showed a significant interaction between habitat and species (t = -5.027, p<0.0001; Figure 4; Table S3). In rural habitats, *C. versicolor* was found on substrates with significantly lower temperatures compared to *P. dorsalis* (t = -5.76, p < 0.0001). Substrate temperature did not differ between the species in the urban habitat (t = 1.64, p = 0.1). Within species, *P. dorsalis* were found on substrates with significantly higher temperatures in rural than urban habitats (t = 6.35, p < 0.0001), while *C. versicolor* showed no significant difference between the habitats (t = -2.29, p = 0.02).

All four populations showed a strong positive correlation between substrate temperature and body temperature, with the degree of thermoregulation (k) being highest for urban populations of *C. versicolor (*R ^2^ = 0.62, k = 0.68) and *P. dorsalis*(R ^2^= 0.620, k = 0.67), followed by rural populations of *C. versicolor (*R ^2^= 0.4, k = 0.47) and *P. dorsalis*(R ^2^= 0.37, k = 0.37; Figure 5).

**Figure 5.**
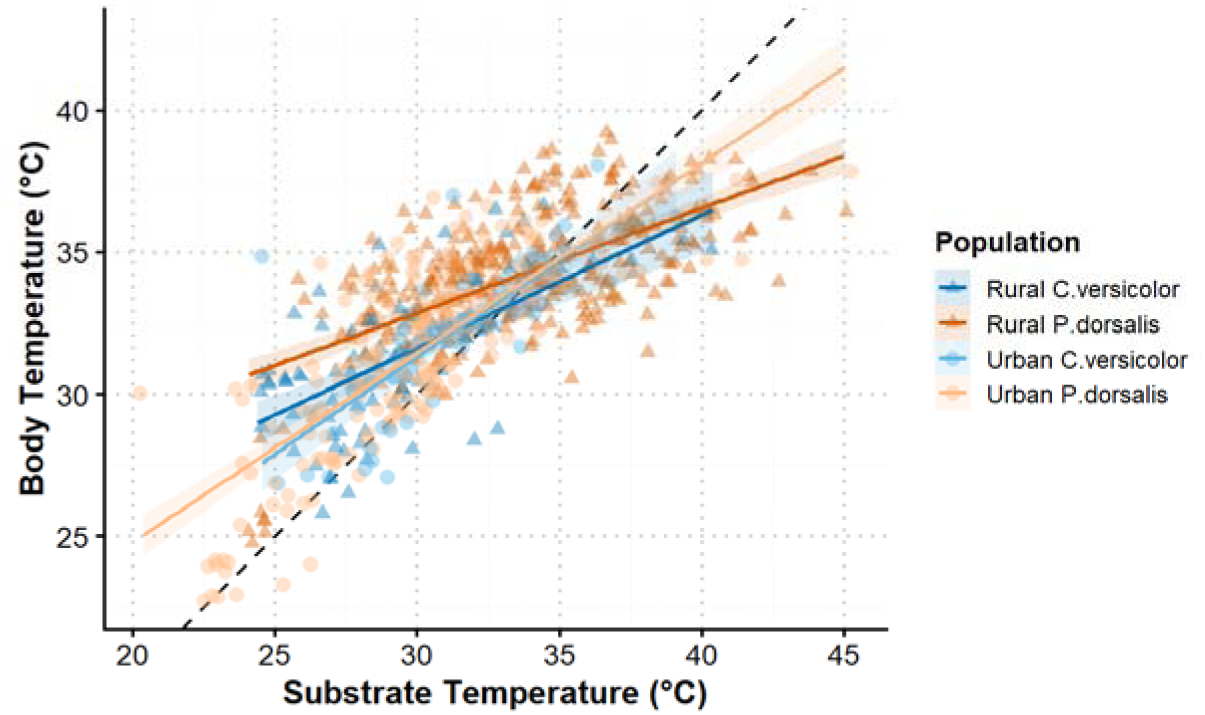
Relationship between body temperature and substrate temperature for *P. dorsalis* and *C. versicolor*, measured from the thermal images of free-ranging lizards from rural and urban environments. Coloured lines denote linear regression slopes and the dashed line denotes a 1:1 relationship.

## 4. DISCUSSION

Urbanization introduces novel environmental conditions, altering both physical structure and thermal properties of habitats. These urban settings can serve as proxies for future climatic conditions, offering insights into how species might respond to changing habitat conditions. However, realistic assessments of species vulnerability to temperature change requires accounting for the thermal heterogeneity and buffering potential of microhabitats. Integrating both species-specific thermal limits and preferences alongside fine scale microclimate data can significantly improve the accuracy of such models (Scheffers et al., 2014). Using Infrared Thermography (IRT) and laboratory trials, we find clear differences in the thermal environment and thermal traits of two co-occurring lizard species across rural and urban environments. Despite species specific differences in microhabitat preferences, urban habitats were similar in terms of thermal heterogeneity and mean substrate temperature used by both species, while the same was not true for rural ones. Moreover, urban habitats favored thermoconformity in both species, as expected for environments where costs and opportunity for thermoregulation are high (Huey & Slatkin, 1976). Within species, *P. dorsalis* experiences warmer and more thermally heterogeneous conditions in the rural than urban habitats, while microhabitat temperatures and thermal heterogeneity were similar across habitat types for *C. versicolor*. Among the thermal traits measured, *C. versicolor* exhibited a broader thermal tolerance (higher CT_max_ and lower CT_min_) and preferred temperature (T_pref_) range than *P. dorsalis*, across both habitats. Overall, our results indicate that thermal responses to urbanization are shaped by species specific thermal traits and patterns of microhabitat use.

Ectotherms show three major responses to microhabitat thermal conditions when achieving their thermal preference (Cabon et al., 2023). Firstly, an adaptation of local optima is observed when individuals actively prefer temperatures (T_pref_) that match conditions experienced in their natural habitat, in a co-gradient manner (Brattstrom, 1965; Keller & Seehausen, 2012). Secondly, a counter-gradient response is observed when individuals from harsher environments show greater thermal sensitivity and select temperatures that negatively correlate with ambient temperatures in their habitat (Fangue et al., 2009; Kuyucu & Çağlar, 2016). And thirdly, a Bogert effect, where behavioural thermoregulation allows individuals to buffer against environmental extremes, resulting in no correlation between measured climatic conditions and T_pref_ (Bogert, 1949; Farallo et al., 2018; Muñoz, 2022). Our results indicate different thermoregulatory patterns in *P. dorsalis* and *C. versicolor* across rural and urban environments. Although urban microhabitats were relatively cooler, urban individuals of *P. dorsalis* preferred higher body temperature than their rural conspecifics in a counter gradient manner. Such patterns can arise when individuals select higher temperatures to compensate for potential lower opportunities to find favourable temperatures in cooler or more constrained environments (Hodgson & Schwanz, 2019; Llewelyn et al., 2017; Matthew T. McElroy, 2014). In contrast, *C. versicolor*, displayed a co-gradient pattern, with rural habitats being warmer and individuals from these habitats showing higher T_pref_ compared to their urban counterparts. This adaptation to local thermal optima suggests that *C. versicolor* may adjust its thermal preferences in response to local climate regimes to maintain optimal physiological performance (Drakulić et al., 2017; Zheng & Liu, 2010). Similar patterns of variation in preferred body temperature, both among (Ćorović et al., 2024; Klinges et al., 2024) and within species (Cabon et al., 2023; Hodgson & Schwanz, 2019) have been reported across a wide range of taxa.

Between species, *C. versicolor* exhibited a significantly broader T_pref_ range than *P. dorsalis* across both habitat types. *Psammophilus dorsalis*, thus shows greater precision in temperature selection, indicating thermal specialization, while *C. versicolor* displayed a more generalist thermoregulatory strategy by exploring a broader range of temperatures along the gradient (Preston & Johnson, 2022). These contrasting strategies reflect fundamental trade-offs associated with thermoregulation that influence species distributions and habitat use (Matt T. McElroy, 2007). Thermal specialists gain significant physiological benefits when maintaining a narrow optimal body temperature but face high energetic costs in doing so, particularly in environments with fluctuating temperatures. Ultimately, this limits their habitat range and geographic distribution. In contrast, thermal generalists function effectively across a wider range of temperatures, significantly extending their potential range and distribution (Huey & Slatkin, 1976). Our findings support this framework, with *C. versicolor* having a widespread geographic range spanning multiple countries across Asia (Radder, 2006), while *P. dorsalis* is mostly restricted to Peninsular India (Daniel, 2002).

Patterns from thermal tolerance further lend support to these conclusions. *Calotes versicolor* showed both a higher upper thermal limit (CT_max_) and a lower thermal limit (CT_min_) and therefore, a wider thermal tolerance breadth compared to *P. dorsalis*. While CT_max_ differed significantly between species, variation across urban and rural populations within each species was minimal. CT_min_, however, showed significant variation both between species and among populations. These findings align with numerous studies which consistently report limited plasticity in upper thermal limits relative to lower thermal limits (Addo-Bediako et al., 2000; Apte et al., 2025). The evolutionary adaptive potential for plasticity in CT_max_ appears to be constrained, especially for tropical ectotherms that already experience temperatures close to their upper thermal limits (Deutsch et al., 2008; Hoffmann et al., 2013). Collectively, broader thermal tolerance breadth and thermal preference observed in *C. versicolor* aligns with its extensive geographic distribution and supports the hypothesis that species with wider thermal niche are more likely to occupy larger geographic areas (Calosi et al., 2008; Kellermann et al., 2009).

*Psammophilus dorsalis* occupied significantly warmer substrates than *C. versicolor* in rural habitats, consistent with their differing microhabitat preferences. However, no significant interspecific differences in substrate temperatures were observed in urban settings. Urban environments may be thermally more homogenized, limiting available microclimate diversity despite species specific differences in habitat use. Such thermal homogenisation likely arises from the spatial configuration of human-made infrastructure and vegetation in urban environments (Pincebourde & Suppo, 2016). Within species, *C. versicolor* experiences similar substrate temperatures across habitats, indicating thermal adaptability, while *P. dorsalis* experiences warmer substrates in rural than urban areas contrary to expectations of the urban heat island effect (Battles & Kolbe, 2019). This observation can be credited to Bengaluru’s urban landscaping which consists of numerous water bodies and abundant green cover, which may provide cooler microhabitat temperatures for urban dwelling lizards (Siddiqui et al., 2021; Thaker et al., 2022).

Thermal heterogeneity within microhabitats, measured using diversity indices, also indicated species and habitat specific patterns. Within species, *P. dorsalis* used areas with significantly higher thermal diversity (SHDI, SIDI) in rural than urban habitats. In contrast, the urban microhabitats used by *C. versicolor* had higher thermal diversity as measured by SHDI while SIDI showed no difference between the habitat types. This suggests that the dominant temperature range was comparable across habitat types and an increased thermal diversity in urban habitats was mostly driven by rarer thermal conditions towards the warmer thermal extremes (Nagendra (2002), Figure 4a). Likewise, for a diurnal gecko species, anthropogenic microhabitats showed more variable microclimates than natural microhabitats, further emphasizing that thermal exposure within microhabitats is highly species specific (Apte et al., 2025). *Psammophilus dorsalis* occupies rocky outcrops with vegetation offering a mosaic of sunlit and shaded microhabitats in rural habitats (Mohanty et al., 2021). These heterogeneous thermal patches facilitate behavioral thermoregulation by allowing individuals to shuttle between microclimates for optimal thermoregulation (e.g., Angilletta et al., 2002; Kearney et al., 2009). However, in an urban setting, the species seem to have limited access to thermally diverse substrates beyond concrete walls ledges. On the other hand, *C. versicolor* is a more generalist and semi arboreal species and therefore may better exploit the different thermal niches in an urban environment provided by structures such as walls, vegetation and shaded areas. Thus, similar thermal heterogeneity between urban and rural habitats may suggest greater thermal flexibility and ability to use behavioural means to buffer against varying thermal conditions.

The model put forth by Huey and Slatkin (1976) predicts that thermoregulatory strategies fall along a continuum from strict thermoregulation to thermoconformity, depending on the costs and benefits imposed by the environment. In extreme environments where precise thermoregulation is energetically costly, individuals are expected to show greater degree of thermoconformity (Angilletta, 2009). We find higher levels of thermoconformity in urban individuals of both species, suggesting a greater cost of thermoregulation. However, thermal heterogeneity and spatial structure of microclimates are as important as mean temperature in determining the thermal quality of habitats (Sears & Angilletta, 2015). Structurally heterogeneous habitats provide diverse microclimates that facilitate effective thermoregulation, while more homogeneous environments may favour thermoconformity and reduced variation in thermal traits (Huey et al., 2003; Sears et al., 2011; Stearns, 1992). Between the two species, urban individuals did not differ markedly in habitat thermal heterogeneity, average substrate temperature used or degree of thermoconformity. However, within the species, urban and rural habitats strongly differed in all these parameters for *P. dorsalis*, but not for *C. versicolor*. These findings imply that *P. dorsalis* is likely more negatively impacted by habitat modification and thus more vulnerable to changes in microclimatic conditions, whereas *C. versicolor* may have a greater capacity to adapt to such changes. Such species-specific differences in thermal physiology and microclimate use are often overlooked in models that assess species vulnerability to changing climate, leading to biased estimation of species risk. For instance, (Sinervo et al., 2010) projected that up to 39% of lizard populations and 20% of lizard species could go extinct worldwide by 2080. However, subsequent works such as (Varner & Dearing, 2014) caution that local microclimatic variation can allow species to persist in landscapes that are predicted to be unsuitable when assessed using large-scale climate data alone.

Overall, our findings demonstrate that physiological responses to urbanization are strongly mediated by species specific thermal physiology and microhabitat use. Both species showed different responses to urbanization, with *P. dorsalis* appearing to be more susceptible to the thermal constraints imposed by human-modified landscapes. These results provide a foundation for future work examining the extent to which local populations can adapt to urbanization, particularly through physiological or behavioral means. Importantly, our study reinforces the growing consensus that maintaining and enhancing thermal heterogeneity within landscapes is critical for the persistence of small ectotherms. Towards this, the conservation and creation of thermally diverse habitats and better urban landscaping would be effective strategies to mitigate the impacts of urbanization and changing temperatures on ectothermic species.

## Supporting information

Supplementary_files_Razak_et_al.2026

## Acknowledgements

Funding for this project was provided by DST-FIST grant to CES, IISc, and by Institute of Eminence (IoE) funding from the Indian Institute of Science. We thank Nishta Rajan, Abhinya P, Harrington Deva and Siddharth Tripathi for their help in carrying out fieldwork. We sincerely thank all members of Thaker lab for their feedback and suggestions during the course of the work.

## Author Contributions

M.R contributed to conceptualisation, data curation, formal analysis, project administration, investigation, methodology, software development, visualisation, writing of the original draft, and review and editing. A.B contributed to conceptualisation, data curation, investigation, methodology, reviewing and project administration. S.D contributed to data curation, investigation, software development, reviewing and visualisation. M.T contributed to conceptualisation, funding acquisition, resources, project administration, supervision, and review and editing.

## Disclosure Statements

### Conflict of Interest

The authors declare no conflicts of interest. All co-authors have seen and agree with the contents of the manuscript and there is no financial interest to report.

### Ethical Guidelines

The ethical approval for this study was obtained from the Institutional Animal Ethics Committee of the Indian Institute of Science (CAF/Ethics/739/2020). All applicable international, national, and institutional guidelines for the use of animals were followed.

### Artificial Intelligence Generated Content

AI-based tools were used for language editing only. Authors are fully responsible for all content.

## Notes

### Competing Interest Statement

The authors have declared no competing interest.

