## Supplementary_files_Razak_et_al.2026 for "Can tropical agamid lizards physiologically tolerate the altered thermal mosaic of urban microhabitats?"

### FIGURES

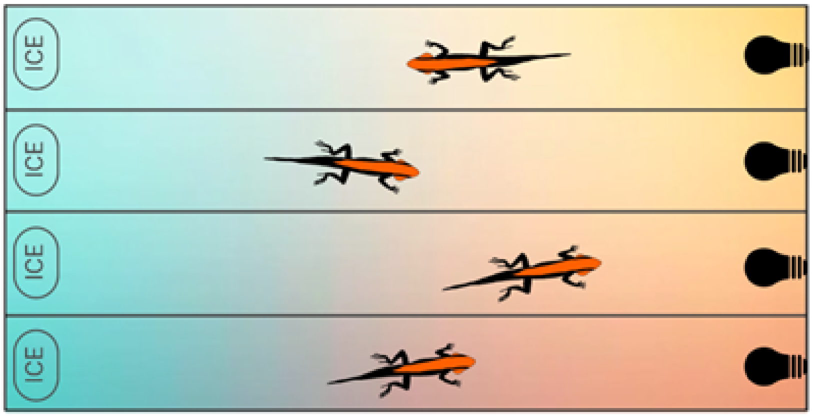


Figure S1

Figure S1. An illustrative diagram of *Psammophilus dorsalis* in the thermal gradient used for Tpref experiments. A temperature gradient is maintained by ceramic infrared heat lamps (hot) on one end and frozen ice packs (cold) on the other.

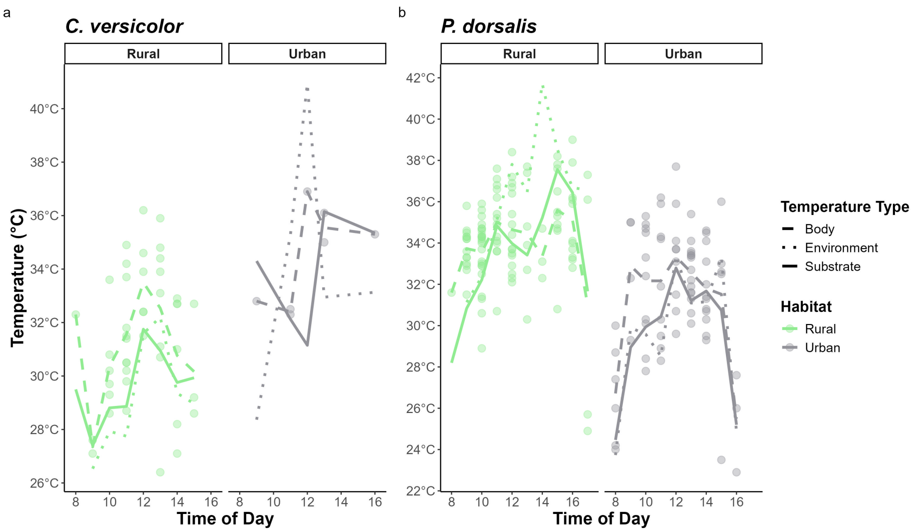


Figure S2

Figure S2 . Variation in mean body temperature, substrate temperature, and environment temperature as measured from the thermal images across the day for *C. versicolor* and *P. dorsalis*. The solid line represents the mean body temperature, the dashed line indicates the substrate temperature, and the dotted line represents the environmental temperature.

Table S1: Thermal preference (Tpref), critical thermal maximum (CTmax), and critical thermal minimum (CTmin) for rural and urban populations of *C. versicolor* and *P. dorsalis* . Values are presented as median ± SD with interquartile range (25–75%). Sample size for each of the measured trait is represented as *N*

| | *C. versicolor* | | | | *P. dorsalis* | | | |
|---|---|---|---|---|---|---|---|---|
| | Rural | | Urban | | Rural | | Urban | |
| | Male | Female | Male | Female | Male | Female | Male | Female |
| Tpref | | | | | | | | |
| *N* | 24 | 2 | 21 | 2 | 38 | 30 | 32 | 19 |
| | 28.23 ± 3.57 | 27.25 ± 7.71 | 30.50 ± 5.23 | 31.48 ± 6.89 | 25.35 ± 2.17 | 25.60 ± 1.70 | 27.70 ± 4.14 | 30.20 ± 3.35 |
| | 26.21–30.06 | 24.53–29.98 | 27.60–34.30 | 29.04–33.91 | 24.63–26.68 | 24.23–26.30 | 26.68–28.63 | 27.65–32.80 |
| CTmax | | | | | | | | |
| *N* | 24 | 2 | 20 | 2 | 38 | 29 | 33 | 19 |
| | 45.45 ± 1.69 | 44.75 ± 1.06 | 46.10 ± 1.31 | 43.30 ± 1.41 | 42.00 ± 2.04 | 40.50 ± 1.98 | 42.90 ± 2.02 | 42.00 ± 2.06 |
| | 44.68–46.33 | 44.38–45.13 | 44.90–46.85 | 42.80–43.80 | 40.40–43.20 | 39.50–42.30 | 40.00–43.50 | 40.35–42.85 |
| CTmin | | | | | | | | |
| *N* | 22 | 2 | 21 | 1 | 37 | 29 | 33 | 20 |
| | 13.85 ± 1.30 | 15.80 ± 5.66 | 13.80 ± 1.77 | 15.90 | 17.20 ± 2.36 | 18.00 ± 2.75 | 19.30 ± 0.73 | 17.50 ± 3.78 |
| | 13.13–14.90 | 13.80–17.80 | 12.50–14.80 | 15.90 | 14.90–18.60 | 16.60–19.00 | 18.60–19.60 | 12.55–19.53 |

RAZAK ET AL.

RAZAK ET AL.



Table S2: Thermal heterogeneity indices measured from thermal images, presented as SHDI (Shannon Diversity Index) and SIDI (Simpson Diversity Index) for urban and rural populations of *P. dorsalis* and *C. versicolor*. Values are presented as median ± SD and interquartile range (25–75%). Bold values denote statistical significance at $p < 0.05$.

| | *C. versicolor* | | *P. dorsalis* | | Statistical Results | | |
|---|---|---|---|---|---|---|---|
| | Rural | Urban | Rural | Urban | Species | Habitat | Interaction Species*Habitat |
| *n* | 120 | 21 | 184 | 202 | | | |
| SHDI | 2.37 ± 0.50<br>2.17–2.73 | 3.00 ± 0.44<br>2.79–3.30 | 2.87 ± 0.46<br>2.59–3.19 | 2.03 ± 0.74<br>1.60–2.60 | $t = 5.82$<br>**p < 0.001** | $t = 3.67$<br>**p < 0.001** | $t = -8.45$<br>**p < 0.001** |
| SIDI | 0.88 ± 0.08<br>0.85–0.92 | 0.92 ± 0.04<br>0.91–0.95 | 0.93 ± 0.04<br>0.91–0.95 | 0.84 ± 0.15<br>0.73–0.91 | $t = 3.80$<br>**p < 0.001** | $t = 1.96$<br>**p = 0.05** | $t = -6.38$<br>**p < 0.001** |

Table S3: Mean body temperature and substrate temperature in urban and rural populations of *P. dorsalis* and *C. versicolor*, from thermal images of lizards in the wild. Values are presented as median ± SD and interquartile range (25–75%). Asterisks (*) denote statistical significance.

| | *P. dorsalis* | | *C. versicolor* | | Statistical Results | | |
|---|---|---|---|---|---|---|---|
| | Urban | Rural | Urban | Rural | Species | Habitat | Interaction Species*Habitat |
| *N* | 146 | 206 | 44 | 56 | | | |
| Body Temperature | 32.00 ± 3.17 | 34.30 ± 2.30 | 32.75 ± 2.66 | 31.25 ± 2.52 | $t = 4.98$ | $t = 1.39$ | $t = -4.27$ |
| | 30.00–34.10 | 33.10–35.55 | 30.97–34.42 | 29.67–32.90 | **p < 0.001** | $p = 0.17$ | **p < 0.001** |
| Substrate Temperature | 30.15 ± 3.60 | 33.80 ± 3.75 | 31.62 ± 2.79 | 28.95 ± 3.46 | $t = 5.76$ | $t = 2.29$ | $t = -5.03$ |
| | 28.30–31.70 | 31.30–36.00 | 29.60–33.82 | 27.05–31.25 | **p < 0.0001** | **p = 0.022** | **p < 0.0001** |

RAZAK ET AL.
